# Dissemination of IncI plasmid encoding *bla*_CTX-M-1_ is not hampered by its fitness cost in the pig’s gut

**DOI:** 10.1101/2023.02.03.527097

**Authors:** Margaux Allain, Anne Claire Mahérault, Benoit Gachet, Caroline Martinez, Bénédicte Condamine, Mélanie Magnan, Isabelle Kempf, Erick Denamur, Luce Landraud

## Abstract

Multiresistance plasmids belonging to the IncI incompatibility group have become one of the most pervasive plasmid types in extended-spectrum beta-lactamase producing *Escherichia coli* of animal origin. The extent of the burden imposed on the bacterial cell by these plasmids seems to contribute to the emergence of “epidemic” plasmids. However, *in vivo* data in the natural environment of the strain are scarce. Here, we investigated the cost of a *bla*_CTX-M-1_-IncI1 epidemic plasmid in a commensal *E. coli* animal strain, UB12-RC, before and after oral inoculation of fifteen 6-to 8-week-old specific pathogen-free pigs. Growth rate in rich medium was determined on (i) UB12-RC and derivatives, with or without plasmid, *in vivo* and/or *in vitro* evolved, and (ii) strains that acquired the plasmid in the gut during the experiment. Although *bla*_CTX-M-1_-IncI1 plasmid imposed no measurable burden on the recipient strain after conjugation and during the longitudinal carriage in the pig’s gut, we observed a significant difference in the bacterial growth rate between IncI1 plasmid-carrying and plasmid-free isolates collected during *in vivo* carriage. Only a few mutations on the chromosome of the UB12-RC derivatives were detected by whole-genome sequencing. RNA-Seq analysis of a selected set of these strains showed that transcriptional responses to the *bla*_CTX-M-1_-IncI1 acquisition were limited, affecting metabolism, stress response, and motility functions. Our data suggest that the effect of IncI plasmid on host cells is limited, fitness cost being insufficient to act as a barrier to IncI plasmid spread among natural population of *E. coli* in the gut niche.

## INTRODUCTION

Since the 2000s, a global epidemic of Extended Spectrum Beta-Lactamase (ESBL)-producing *Enterobacterales* has emerged, linked to the dissemination of successful *bla*_CTX-M_-carrying clones (1). The CTX-M enzymes are mostly plasmid encoded and *E. coli* is the major host (2).

Among major multiresistant plasmids in *Enterobacterales*, IncF, IncI, IncC, IncN, and IncL plasmid families are the most commonly reported. The I-complex plasmids contain incompatibility groups I, K, B, and Z. IncI, or MOB_P_ according to relaxase typing, is a group of low copy-number, narrow-host-range, conjugative plasmids, which vary in size from 50 to 250 kb (2, 3). Some indistinguishable IncI1 and IncIγ plasmids have been isolated from multiple, unrelated high-risk clones, identified in different periods and bacterial species, in distant geographical areas, defined as “epidemic” plasmids (4–7). IncI pST3 plasmid is the most commonly reported *bla*_CTX-M-1_-IncI1 plasmid in Europe, described in pathogenic human *E. coli* isolates, and in *E. coli*, and *Salmonella spp*. from food-producing animals (8). Moreover, Lucas *et al*. recently showed that resistance to Extended-Spectrum Cephalosporins (ESC) in French pigs’ *E. coli* isolates is mainly carried by highly similar *bla*_CTX-M-1_-IncI1/ST3 plasmids (9). IncI plasmids have been recognized in clinically relevant bacteria from human, animal, and environmental sources, and as major vehicles for the dissemination of ESBL, notably CTX-M-15 and CTX-M-1 enzymes (2, 8).

Previous studies using a series of experiments that included pigs inoculated with *E. coli* harboring *bla*_CTX-M-1_-encoding IncI plasmid showed the maintenance of the plasmid in its *E. coli* host, in the pig’s gut. Moreover, the loss of the plasmid was a rare event in gut microbiota during these longitudinal studies of *in vivo* carriage in pigs (10). In these models, the transfer of the *bla*_CTX-M-1_-encoding IncI plasmid between different *E. coli* strains occurred and underlines the dissemination capacity of the ESC resistance, independently of the exposure to antibiotic drugs (11, 12). Finally, understanding the factors of IncI1 plasmids success better is important, as they are major contributors to the dissemination of ESBL genes especially in the animal reservoir.

The arrival of a plasmid in a new bacterium produces a highly variable fitness cost depending on the plasmid–bacterium association (13–15). Plasmid persistence and dissemination in bacterial population, under conditions that do not select for plasmid-encoded genes, are influenced by the fitness effects associated with plasmid carriage (16, 17). Previous works have shown that compensatory adaptation frequently ameliorates the fitness of costly plasmids (18–20). This compensatory evolution may play a key role in the dynamics of the spread and evolution of antibiotic resistance plasmids (13, 21–23). However, most studies on the evolutionary dynamics of plasmid–bacterium associations have been performed in laboratory-based culture systems.

We took the unique opportunity of the previous studies performed in pigs for probiotic testing (12), to evaluate the fitness cost of IncI pST3 plasmid on bacterial isolates collected during an *in vivo* longitudinal study of carriage of a CTX-M-1 producing *E. coli* B1 commensal strain in pigs. We also investigated the impact of IncI pST3 plasmid on the bacterial host using whole-genome sequencing (WGS) and transcriptome sequencing and analysis to determine the effect of plasmid carriage on global gene expression.

## RESULTS

### Characterization of ancestral and *in vivo* evolved UB12 strains

According to the whole-genome sequencing results, *E. coli* UB12-RC recipient strain belongs to O100:H40 serotype, B1 phylogroup, and ST2520, according to the Warwick multilocus sequence typing (MLST) scheme. This strain harbors a mutation in *rpoB* after selection in Mueller-Hinton medium (MH) containing 250 mg/liter rifampicin (10), and contains two plasmids: one P1-like phage-plasmid (IncY, 98119 bp) (24), and one small mobilizable plasmid (Col156 plasmid, 5223 bp) (Table 1). In addition to the IncY and Col156 resident plasmids, the UB12-TC transconjugant harbors a *bla*_CTX-M-1_-IncI plasmid (108699 bp), and one colicinogenic plasmid, belonging to the ColE1 group (ColRNAI plasmid, 6724 bp), which disseminated from the M63-DN donor strain. The *bla*_CTX-M-1_-IncI plasmid is of plasmid multilocus sequence type 3 (pST3) of the IncI1 family and presents 99% identity and 72 % coverage to R64 plasmid, a prototype of the IncI1 group (ascension N° AP005147)(8). Moreover, the core genome of IncI ST3 plasmid presents with other *bla*_CTX-M-1_-IncI1/ST3 plasmid previously described 100% identity and 99% coverage (ascension n° KJ484635)(25). The *bla*_CTX-M-1_ gene is associated with an *IS*Ecp1 mobile element found at the 5’ end of the gene, and the *IS*Ecp1-*bla*_CTX-M-1_ transposition unit is inserted in the shufflon region of the plasmid. Other resistance genes carried in the *bla*_CTX-M-1_-IncI/ST3 plasmid are *dfrA17, sul2*, and *aadA5* (Table 1).

**Table 1:** Characterization of donor *E. coli* strain, evolved and ancestral *E. coli* UB12 clones, and new in vivo transconjugants used in this study.

| status | Isolate name | Experiment (month/year) | Box<sup>a</sup> | Animal<sup>a</sup> | Day of evolution | Phylogroup | MLST type<sup>b</sup> | Serotype | FimH Allele | Plasmid replicon<sup>c</sup> | Antimicrobial resistance genes | Mobility<sup>d</sup> |
| --- | --- | --- | --- | --- | --- | --- | --- | --- | --- | --- | --- | --- |
| Ancestral strains | M63-DN |  | - | - | - | B1 | ST2521 | O147:H7 | fimH31 | **Incl (KJ484635)**; ColRNAI (J01566) | *bla*<sub>CTX-M-1</sub>, *sul*2, *aadA5*, *dfrA17* | + |
|  | UB12-RC |  | - | - | - | B1 | ST2520 | O100:H40 | fimH31 | **IncY (MH42254)**; Col156 (EU090225) |  | + |
|  | UB12-TC |  | - | - | J0 | B1 | ST2520 | O100:H40 | fimH31 | **Incl (KJ484635)**; ColRNAI (J01566); **IncY (MH42254)**; Col156 (EU090225) |  | +/- |
| Evolved strains with Incl plasmid | UB12-TC<sup>IVE</sup>.1 | January/2013 | F3v | 4368 | D28 | B1 | ST2520 | O100:H40 | fimH31 | **Incl (KJ484635)**; ColRNAI (J01566); **IncY (MH42254)**; Col156 (EU090225) | *bla*<sub>CTX-M-1</sub>, *sul*2, *aadA5*, *dfrA17* | +/- |
|  | UB12-TC<sup>IVE</sup>.2 | June/2013 | B2d | 238 | D8 |  |  |  |  |  |  | +/- |
|  | UB12-TC<sup>IVE</sup>.3 | June/2013 | B4v | 4631 | D31 |  |  |  |  |  |  | +/- |
|  | UB12-TC<sup>IVE</sup>.4 | October/2012 | B1 | 4219 | D7 |  |  |  |  |  |  | +/- |
|  | UB12-TC<sup>IVE</sup>.5 | October/2012 | B1 | 4219 | D20 |  |  |  |  |  |  | +/- |
|  | UB12-TC<sup>IVE</sup>.6 | December/2015 | C3d | 462 | D18 |  |  |  |  |  |  | +/- |
|  | UB12-TC<sup>IVE</sup>.7 | December/2015 | C4v | 5652 | D45 |  |  |  |  |  |  | +/- |
| Evolved strains with Incl plasmid loss | UB12-TCApIncl.1 | June/2013 | B2v | 4613 | D20 | B1 | ST2520 | O100:H40 | fimH31 | ColRNAI (J01566); **IncY (MH42254)**; Col156 (EU090225) | - | + |
|  | UB12-TCApIncl.2 | April/2012 | C3v | 3716 | D7 |  |  |  |  |  |  | + |
|  | UB12-TCApIncl.3 | April/2012 | C3v | 3716 | D18 |  |  |  |  |  |  | + |
| New in vivo transconjugants with Incl plasmid | NI.1 | January/2013 | B2v | 4331 | D22 | D | ST13022 | -:H25 | fimH123 | **Incl (KJ484635)**; **IncY (MH42254)**; IncFII (pRSB107) | *bla*<sub>CTX-M-1</sub>, *sul*2, *aadA5*, *dfrA17* | + |
|  | NI.2 | January/2013 | F3v | 4324 | D22 | A | ST48 | O184:H11 | FimH41 | **Incl (KJ484635)**; IncFIC(FII), IncFIB (AP001918); IncFII (pSFO) (AF401292); ColE10 (X01654); ColRNAI (J01566) | *bla*<sub>CTX-M-1</sub>, *sul*2, *aadA5*, *dfrA17*, *sul*1, *tetA*, *tetB* | +/- |
|  | NI.3 | January/2013 | F3v | 4368 | D10 | B1 | ST13023 | -:H7 | fimH68 | **Incl (KJ484635)** | *bla*<sub>CTX-M-1</sub>, *sul*2, *aadA5*, *dfrA17* | + |
|  | NI.4 | June/2013 | B2v | 4591 | D16 | B1 | ST2540 | OgN9:H2 | fimH31 | **Incl (KJ484635)**; ColRNAI (J01566) | *bla*<sub>CTX-M-1</sub>, *sul*2, *aadA5*, *dfrA17* | + |
|  | NI.5 | June/2013 | B2v | 4575 | D13 | G | ST657 | O149:H25 | fimH97 | **Incl (KJ484635)**; IncFII(pRSB107)(AJ851089); ColRNAI (J01566); Col(MG828) | *bla*<sub>CTX-M-1</sub>, *sul*2, *aadA5*, *dfrA17* | + |
|  | NI.6 | June/2013 | B2d | 238 | D8 | C | ST410 | Onove14:H9 | fimH24 | **Incl (KJ484635)**; IncFIC(FII), IncFIB (AP001918), IncFII (pSFO) (AF401292); ColRNAI (J01566) | *bla*<sub>CTX-M-1</sub>, *sul*2, *aadA5*, *dfrA17*, *tetA* | - |
|  | NI.7 | October/2012 | B1 | 4219 | D20 | G | ST117 | O24:H4 | fimH97 | **Incl (KJ484635)**; IncFIC(FII), IncFIB, IncF1A (AP001918), IncFII (pSFO) (AF401292); ColRNAI (J01566); Col(MG828) | *bla*<sub>CTX-M-1</sub>, *sul*2, *aadA5*, *dfrA17* | + |
|  | NI.8 | December/2015 | C4d | 5625 | D4 | B1 | ST767 | O8:H9 | fimH32 | **Incl (KJ484635)**; IncFIC(FII), IncFIB (AP001918); IncFII (pSFO) (AF401292); Col(MG828) | *bla*<sub>CTX-M-1</sub>, *sul*2, *aadA5*, *dfrA17*, *tetA* | + |
|  | NI.9 | December/2015 | C4d | 5625 | D11 | B2 | ST95 | O18:H7 | fimH15 | **Incl (KJ484635)**; IncFIC(FII), IncFIB (AP001918), IncFII (pSFO) (AF401292); ColRNAI (J01566) | *bla*<sub>CTX-M-1</sub>, *sul*2, *aadA5*, *dfrA17* | +/- |
|  | NI.10 | December/2015 | C4v | 5626 | D11 | G | ST117 | -:H19 | fimH97 | **Incl (KJ484635)**; IncFIB (AP001918), IncFII (pRSB107); Col(MG828); Col8282 (DQ995353); p0111; ColRNAI (J01566) | *bla*<sub>CTX-M-1</sub>, *sul*2, *aadA5*, *dfrA17*, *tetB* | - |
|  | NI.11 | December/2015 | C4d | 5651 | D11 | C | ST88 | O8:H9 | fimH39 | **Incl (KJ484635)** | *bla*<sub>CTX-M-1</sub>, *sul*2, *aadA5*, *dfrA17* | -/+ |

<sup>a</sup>, detailed in (9, 11); <sup>b</sup>, according to the Warwick multilocus sequence typing (MLST) scheme; <sup>c</sup>, PlasmidFinder, Incl plasmid is in bold ; <sup>d</sup>, (+) total spreading within medium, (+/-) partial spreading, and (-) not spreading

Three cefotaxime-susceptible and rifampicin-resistant isolates, collected at days 7, 18, and 20, during longitudinal carriage of the UB12-TC strain in the pigs, have spontaneously lost the IncI plasmid *in vivo* and harbor Col156, IncY, and ColRNAI plasmids (referenced as UB12-TCΔpIncI) (Table 1).

To identify mutations and rearrangements that occur during the carriage of UB12-TC strains in the pig’s gut, we used the open-source Breseq pipeline (26) and the ancestral UB12-TC strain as reference genome. In the first step, we excluded all single nucleotide polymorphisms (SNP), insertions or deletions detected comparing mapping reads of the UB12-TC ancestral strain to the assembly of itself. In the second step, we determined genetic differences occurring between the UB12-RC recipient strain, seven UB12-TC isolates, collected in stools between day 7 and 45 after oral inoculation of pigs, i.e. *in vivo* evolved (UB12-TC^ive^), and the cefotaxime-susceptible and rifampicin-resistant isolates (UB12-TCΔpIncI) (Table 2). All transconjugants showed one intergenic point mutation, upstream of the *nadA* gene, except two (Table 2). In chromosomal DNA, we detected seven other events between the UB12-RC recipient strain, the UB12-TC strain and six UB12-TC^ive^ transconjugants: one intergenic point mutation between the *ygeH* and *ygeG* genes, one synonymous point mutation in the *kdsB* gene, and five nonsynonymous point mutations in the *ecpC, ydeE, ttdB, thiH* genes, and in the *hlyE* pseudogene (Table 2). In total, one to three point mutations per strain were detected. In plasmid DNA, a common rearrangement was detected in the shufflon region of the IncI plasmid in three transconjugants. None of other plasmids showed any modification.

**Table 2:**
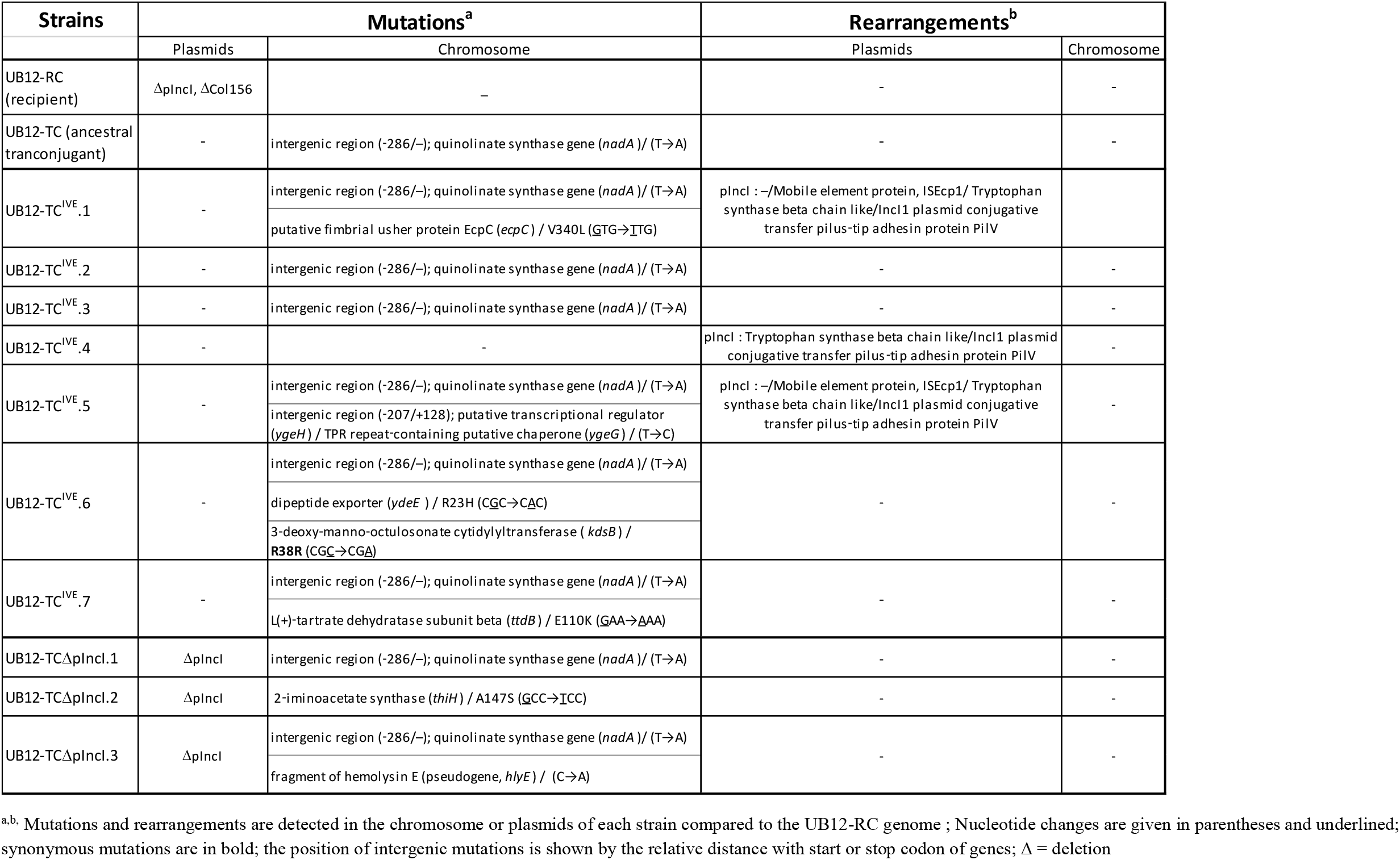
Characterization of mutations and rearrangements in the chromosome and plasmids of ancestral and evolved *E. coli* UB12 clones.

### No fitness cost decrease of UB12 strains during digestive carriage in pigs

To evaluate the IncI plasmid cost on the bacterial fitness, we compared the maximum growth rates (MGRs) of the *E. coli* M63-DN donor strain, the UB12-RC recipient strain and the ancestral UB12-TC transconjugant. The MGR of M63-DN donor strain (3.55 div h-1) was significantly higher than the MGR of UB12-RC strain (3.38 div h-1) (*P*=0.02). The difference between the MGR of UB12-RC strain and the MGR of UB12-TC transconjugant (3.28 div h-1) was nonsignificant (*P*=0.09) (Fig 1).

**Fig 1:**
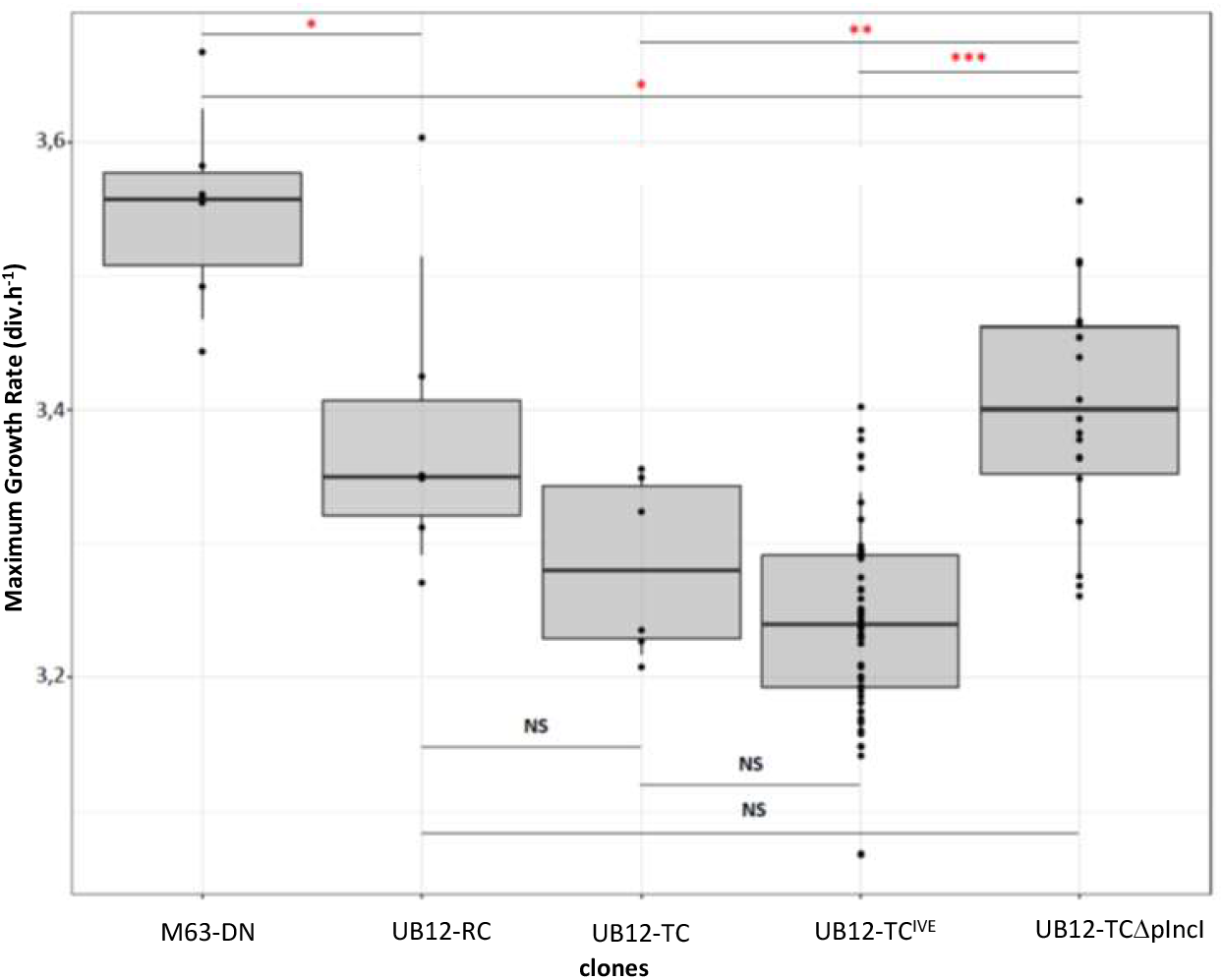
Boxplot representation of the maximum growth rate (MGR) of each pool lineage: MGR of the donor strain (M63-DN), the recipient strain (UB12-RC), the ancestral transconjugant (UB12-TC, IncI, pST3), seven *in vivo* evolved transconjugants (UB12-TC^ive^), and three evolved transconjugants which had lost *in vivo* the IncI plasmid (UB12-TCΔpIncI). UB12-TC^ive^ and UB12-TCΔpIncI were isolated in stools between day 7 and day 45 during longitudinal study of *in vivo* carriage in pigs. For each box, the central mark indicates the median, and the bottom and top edges of the box indicate the 25th and 75th percentiles, respectively. Each point corresponds to one measurement per clone, which were repeated six times per strains. Asterisks indicate significant differences (* = *P* < 0.05, ** = *P* < 0.01 and *** = *P* < 0.001). NS = not significant.

To evaluate the adaptation of IncI plasmid in the *E. coli* UB12 strain, we compared the MGRs of the UB12-TC strain, the M63-DN donor, and ten clones (UB12-TC^ive^ and UB12-TCΔpIncI) isolated from the stools of the pigs, between day 7 and day 45, during an *in vivo* longitudinal study of carriage of the UB12-TC transconjugant (11). Among these clones, three UB12-TC transconjugants had spontaneously lost the IncI plasmid (UB12-TCΔpIncI). As no significant fitness cost variation was detected between the MGRs of the evolved UB12-TC^ive^ transconjugants on the one hand, and between the MGRs of the UB12-TCΔpIncI transconjugants on the other hand, further statistical analyses were performed after pooling the MGRs of the clones from each group. The difference between the MGRs of UB12-TC transconjugants (3.28 div h-1) and the MGRs of UB12-TC^ive^ evolved transconjugants (3.23 div h-1) was nonsignificant (*P*=0.08), as the difference between the MGRs of UB12-RC clones and the MGRs of the UB12-TC^ive^ transconjugants (*P*=0.07). Interestingly, the differences between the MGRs of UB12-TCΔpIncI (3.40 div h-1) and the MGRs of UB12-TC or UB12-TC^ive^ clones were significant (*P*=0.004 and *P*=1.61 10-^7^ respectively) (Fig 1).

### No fitness cost decrease during *in vitro* experiment evolution of UB12-TC

To assess if the absence of decrease in *in vivo* fitness cost could be reproduced *in vitro* in a constant well-defined environment, we conducted an *in vitro* evolution experiment in LB during 30 days (300 generations in total) and evaluated the impact of the IncI plasmid on the bacterial host after evolution. We measured the MGRs of five independent ancestral transconjugants (UB12-TC), and of five independent clones for each lineage after *in vitro* evolution (UB12-TC^ite^), compared to the MGRs of five plasmid-free ancestral and evolved control strains (UB12-RC and UB12-RC^ite^). First, no significant fitness cost variation was detected between the MGRs of the evolved clones belonging to the same lineage. Further statistical analyses were performed after pooling the MGRs of the clones from each lineage. The differences between the MGRs of UB12-TC (3.49 div h-1) and UB12-TC^ite^ (3.35 div h-1) clones, and the MGRs of plasmid-free control strains UB12-RC and UB12-RC^ite^ (3.49 div h-1 and 3.42 div h-1, respectively) were nonsignificant (Fig 2). Although no significant difference was found between ancestral and evolved strains, the MGRs of UB12-TC decreased after the evolution experiment (3.49 div h-1 versus 3.35 div h-1 for UB12-TC and UB12-TC^ite^, respectively). Of note, there was a trend (*P*= 0.054) for increased cost fitness in evolved strains (Fig 2).

**Fig 2:**
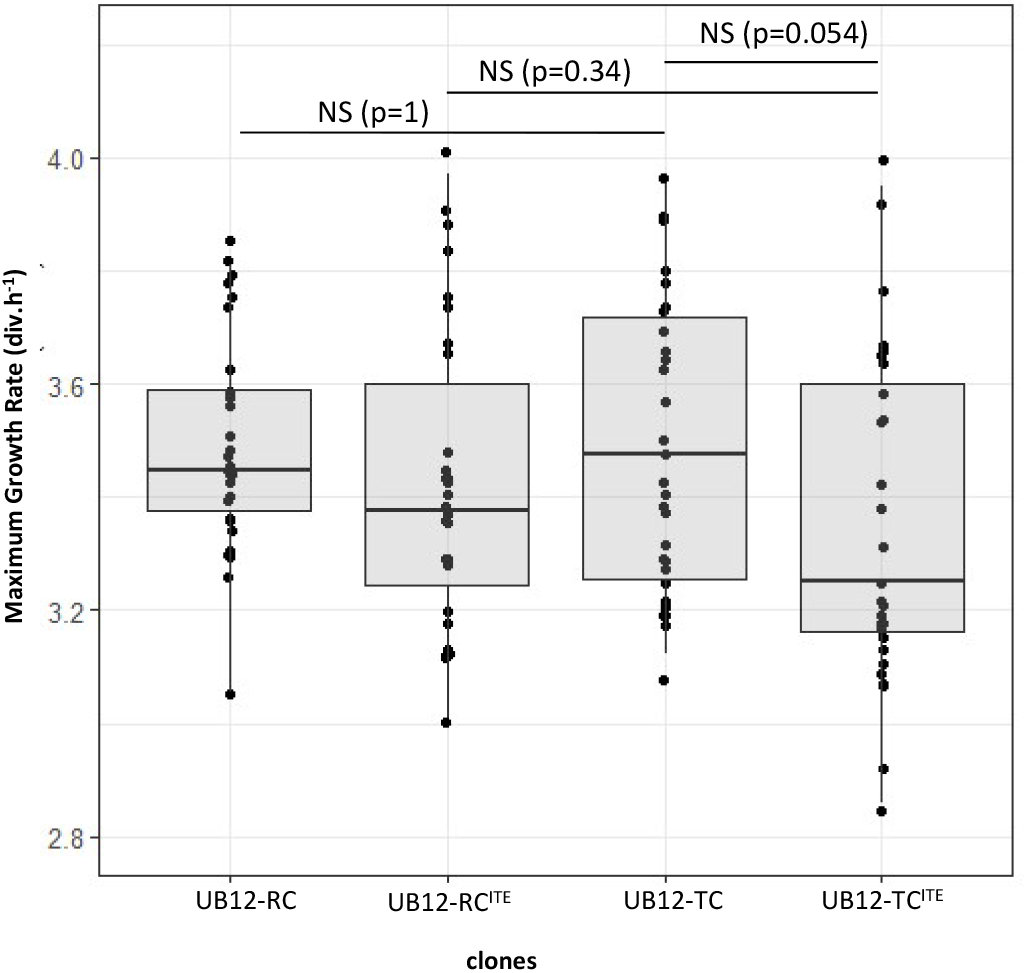
Boxplot representation of the maximum growth rate (MGR) of each pool lineage: MGR of five ancestral UB12-RC and UB12-TC control strains and evolved UB12-RC and UB12-TC strains at day 30. For each box, the central mark indicates the median, and the bottom and top edges of the box indicate the 25th and 75th percentiles, respectively. Each point corresponds to one measurement per clone, which were repeated six times per transconjugant and control. NS = not significant.

### Description of the newly transconjugants *E. coli* isolates (NI) with IncI plasmid

As the recipient host background could interfere in the spread of multidrug resistance plasmids, we analyzed the genetic diversity of eleven cefotaxime-resistant and rifampicin-susceptible newly formed transconjugants (natural isolates, NI) harboring IncI. The phylogroups of NI are diverse (one A, B2, and D, two C, three B1, and G) (Fig 3A). Based on plasmid-families replicon, the plasmidome of NI appears varied with multiple combinations of plasmid types. Most of the eleven isolates (eight NI) carry Col-like plasmids, alone or co-carried with IncF plasmids. One isolate (NI-1) carries an IncY plasmid, sharing 98% identity and 71% coverage to the IncY plasmid of UB12-TC (Table 1). Eight NI (72%) harbor more than two plasmids, while two NI (NI-3 and NI-11) harbor the IncI plasmid alone (Table 1).

**Fig 3:**
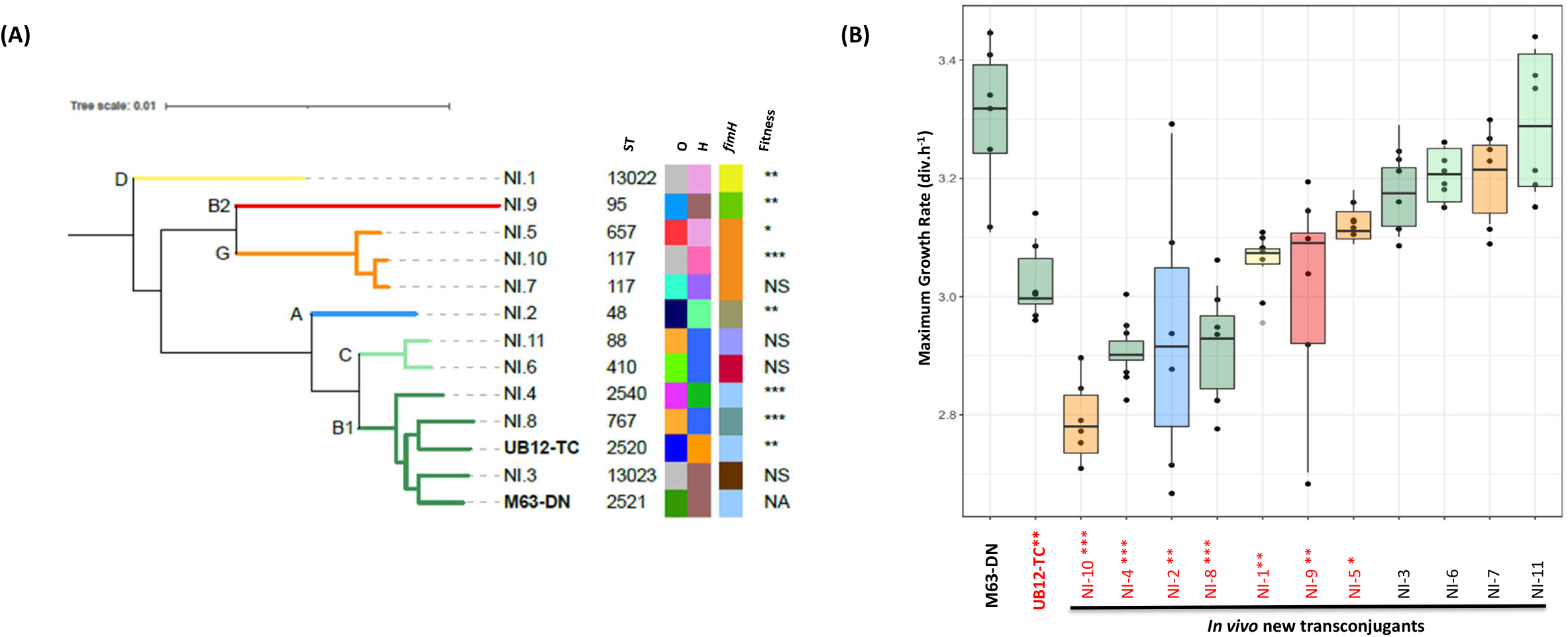
**(A)**, Phylogenetic tree of the 11 newly formed transconjugants detected among pig *E. coli* commensal isolates (NI clones, *E. coli*, IncI, pST3), the M63-DN donor strain, and the UB12-TC transconjugant strain (in bold) are presented. The tree is rooted on an *Escherichia* clade (E1492cladeI, (56)). The phylogenetic tree was constructed with IQ-TREE v1.6.9 from the core genome genes using Roary v1.007002. From left to right are presented the Sequence Type (according to the Warwick University scheme), the O and H serotypes, the *fimH* allele, and the fitness cost of NI clones compared to the donor strain (M63-DN) (NA= not applicable, NS= not significant, * = *P* < 0.05, ** = *P* < 0.01 and *** = *P* < 0.001). Each branch of the same phylogenetic group are colored (yellow, D; red, B2; orange, G; blue, A; light green, C; dark green, B1). Information on O group, H type, and *fimH* allele are given in Table 1. **(B)**, Boxplot representation of the maximum growth rate (MGR) of each strain : donor strain (M63-DN), UB12-TC ancestral transconjugant and NI clones, isolated in stools between day 4 and day 22 during longitudinal study of *in vivo* carriage in pigs. For each box, the central mark indicates the median, and the bottom and top edges of the box indicate the 25th and 75th percentiles, respectively. Each point corresponds to one measurement per clone, which were repeated six times per strains. Asterisks in red indicate significant differences (* = *P* < 0.05, ** = *P* < 0.01 and *** = *P* < 0.001) in comparison to M63-DN. Colors correspond to the strains of the same phylogroup (color code as in A)

The core genomic divergence, estimated by the number of core genome SNPs between strains, is variable and dependent on the sequence type and phylogroup of strains, from 18513 to 77175 (between *E. coli* B1 ST2520 UB12-TC strain, and respectively, *E. coli* B1 ST2540 NI-4 clone and *E. coli* B2 ST95 NI-9 clone) (Fig 3A, Table S1).

### Variable cost of IncI plasmid in transconjugants *E. coli* NI from the pig’s gut

To evaluate the impact of the IncI pST3 plasmid on new bacterial host, we proposed to compare the MGRs of the UB12-TC strain, the M63-DN donor, and eleven cefotaxime-resistant and rifampicin-susceptible NI from the stools of the pigs, between day 4 and day 22. These isolates all harbored the IncI pST3plasmid that have disseminated between different *E. coli* in the pigs’ microbiota after inoculation (Table 1). The MGRs of four NI were not significantly different from that of the donor strain M63-DN, while seven NI showed an MGR significantly lower than those of the M63-DN strain (*P* between 2.7 10^−5^ and 0.015) (Fig 3B).

In order to determine the relative abundance of the newly formed transconjugants in the pig’s gut, we compared the titer of cefotaxime-resistant and rifampicin-susceptible *E. coli* clones to all cefotaxime-resistant *E. coli* population, on the days during which each NI was collected. We observed a variable relative abundance, ranging from 10% to 30% for NI-1, NI-2, NI-3, and NI-6, from 40% to 50% for NI-4, NI-9, NI-10, and NI-11, and more than 65% for NI-5, NI-7, and NI-8.

### Transcriptomic analysis of UB12 strains

We compared the transcriptome of the ancestral UB12-TC transconjugant with the UB12-RC recipient strain to determine the impact of plasmid acquisition on host gene expression. We also compared the transcriptome of two UB12-TC^ive^ transconjugants, isolated between day 28 and 30 in the pig’s gut, and of one UB12-TCΔpIncI transconjugant with the UB12-RC recipient and the ancestral UB12-TC strains to determine the impact of the IncI pST3plasmid maintenance on the bacterial host in the pig’s gut. We combined the results from differential gene expression analysis of two independent biological replicates for each clone, except for UB12-TC.

We observed a low significant transcriptional response of the acquisition of plasmids into UB12-TC, with ten significantly (> 2-log change) differentially expressed genes (adjusted P value (P_adj_) < 0.05) between UB12-RC and UB12-TC strains. Six genes were down-regulated and four genes were up-regulated in UB12-TC compared to UB12-RC. Functions affected by plasmids acquisition included genes involved in mobility and chemotaxis (down-regulation of *flgB* gene; up-regulation of *motA* and *cheA* genes), in stress response (down-regulation of *cyuP* and *ydcI* genes; up-regulation of *gnsA* gene) or genes encoding for phage proteins (down-regulation of three chromosomal genes; up-regulation of one IncY plasmid-gene) (Table 3). No significant change in chromosomal and plasmid gene expression was observed in the UB12-TC^ive^ clones, compared to UB12-TC before carriage in the pig’s gut.

**Table 3:**
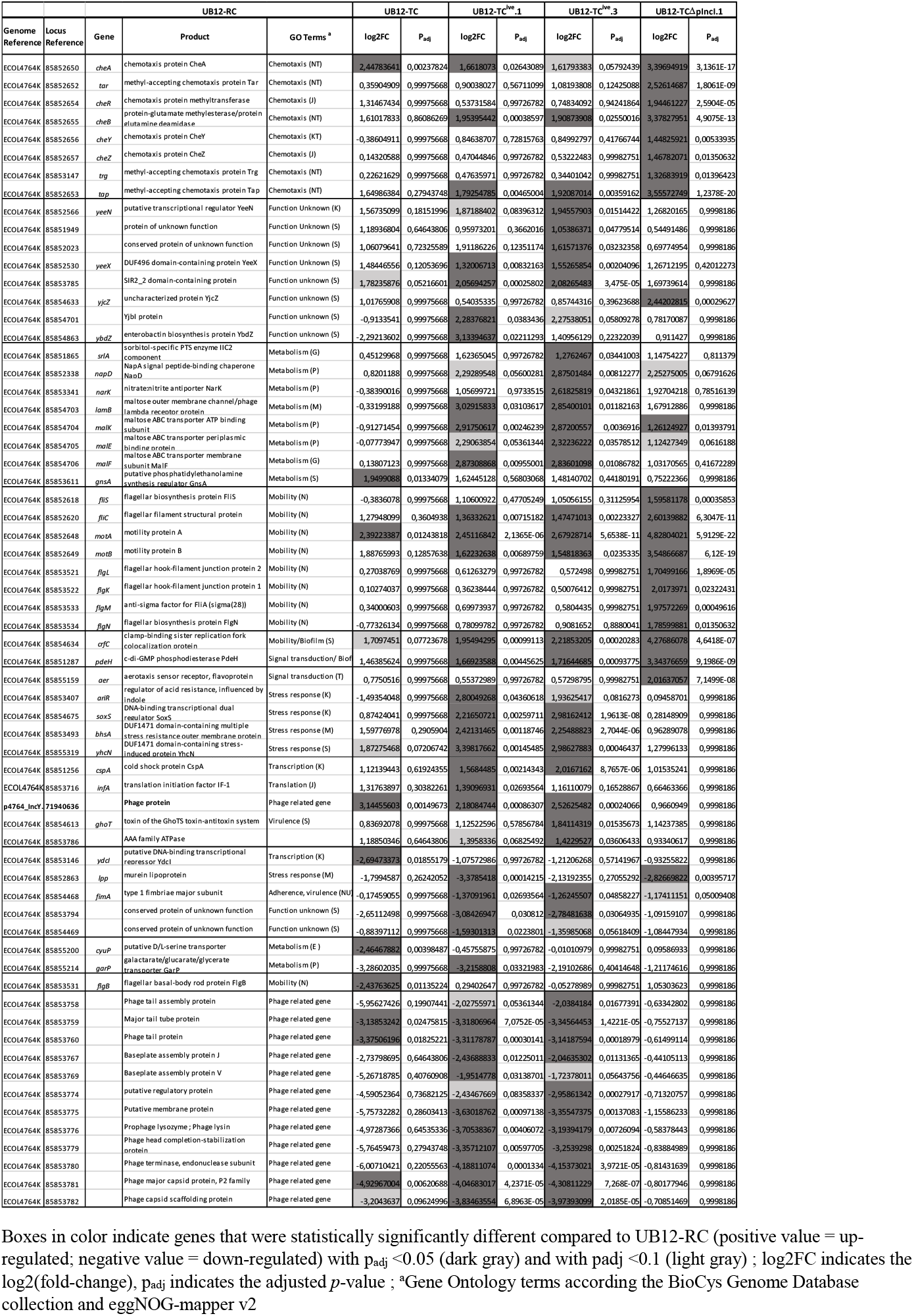
List of differentially expressed genes from UB12-TC clones compared to UB12-RC recipient strain.

However, we detected a transcriptional response between the UB12-RC and UB12-TC^ive^ strains, with a total number of significantly (> 2-log change) differentially expressed genes (P_adj_ < 0.05) ranging from 37 to 39. Among these differentially expressed genes observed between UB12-RC and both UB12-TC^ive^ clones, most of down-regulated chromosomal genes encoded for phage proteins (12 genes). Other down-regulated genes were *fimA* gene, encoding for type 1 fimbriae major subunit, and two genes involved in metabolism (*lpp* and *garP* genes, significantly detected down-regulated only for one of UB12-TC^ive^ clones) (Table 3). In one or both of UB12-TC^ive^ clones, we detected up-regulated genes involved in various stress response, such as *ariR* (oxidative stress), *bhsA* (multiple stress resistance), *yhcN* (hydrogene peroxide stress), c*spA* (temperature response), and s*oxS* (superoxide stress mastor regulator), and genes encoding for transporters in various metabolic pathways (*srlA*, for sorbitol; *narK* or *napD*, for nitrate/nitrite; and *malEFK*, for maltose). Other functional classes of genes detected as significantly up-regulated were those involved in biofilm (*crfC* and *pdeH* genes), in mobility (*motAB* and *fliC* genes), and in chemotaxis (*cheB* and *tap* genes) (Table 3).

All genes involved in mobility and chemotaxis, differentially expressed between UB12-TC^ive^ and UB12-RC strains, were also up-regulated in the UB12-TCΔpIncI clone. Moreover, other genes included in the chemotaxis system (*tar, cheZYR*, and *tgr* genes) and the flagella systems (*fliS, flgN, flgKL*, and *flgM* genes) were up-regulated between UB12-TCΔpIncI and UB12-RC strains. Interestingly, *fliA*, the other major regulator of the flagella systems, appeared significantly differentially expressed (down-regulation) between UB12-TC and UB12-TCΔpIncI clones.

Using the replicase *repA* gene for comparison, we showed that the transcription of IncI plasmid genes was not different between UB12-TC and UB12-TC^ive^ clones (Table S2). We observed that most of the IncI plasmid genes (111 genes, 89.5%) were expressed at low level or not expressed compared to *repA* transcription. Only 13 of predicted CDS (10%) were transcribed at level between 4- and 35-fold higher than *repA*. The most expressed plasmidic gene was annotated as a conserved protein of unknown function, located just upstream of *oriT* and the *nikA* gene (encoding for a Ribbon-helix-helix relaxosome protein), which was transcribed at level 5-fold higher than *repA*. Two other loci in plasmid core regions were transcribed at level between 5- and 16-fold higher than *repA*, the *excAB* genes and an ORF located just upstream, and the *parBM* genes. Two accessory regions of the IncI plasmid encoding resistance genes were also transcribed at high level, the *IS*Ecp-1-*bla*_CTX-M-1_ cassette and three genes (*glmM, sul2*, and *drfA*) located on an *ISCR2* mobile genetic element (Table S2). We observed that the IncY plasmid genes were also expressed at low level. Only 10% (11 genes) of the coding sequences were transcribed at higher level than *repA*, and all annotated as conserved proteins of unknown function (Table S2).

### Swimming motility of strains

To confirm the transcriptomic data on genes involved in mobility, we tested the swimming ability of all the strains used in our study in semi-solid LB agar tubes. We observed that the swimming ability decreased in UB12-TC strains compared to UB12-RC and M63-DN strains. While the ancestral UB12-TC strain presented a comparable swimming motility phenotype to all UB12-TC^ive^ clones, UB12-TCΔpIncI clones presented an increase in the ability to swim in semi-solid LB agar tubes. Of note, we observed various phenotypes, from non-motile to motile, for all eleven NI pig’s clones, which all harbored IncI plasmid (Table 1).

## DISCUSSION

Multiresistance plasmids are widely distributed across bacteria and one of the main drivers of antibiotic resistance spread. The success of a plasmid-borne resistance is constrained by the fitness cost that the multiresistance plasmid imposes on the bacterial cell. It is not clear whether epidemic *bla*_CTX-M-1_-IncI1 plasmids, one of the most common plasmid type in ESBL producers of animal origin (2, 8), conferred a fitness cost on the host bacterium. Previous studies have shown that the *bla*_CTX-M-1_-IncI1 plasmid imposes no or negligible fitness costs on its *E. coli* host *in vitro* (27) and persists without antimicrobial usage (28). On the other hand, using *in vitro* competition experiments, Freire Martín *et al*. showed that the IncI1 plasmid-cured isolate outcompeted the original plasmid-carrying isolate, or plasmid-complemented strains, suggesting a fitness burden was imposed by carriage of this plasmid (29). These studies were performed using *in vitro* models and/or laboratory-adapted strains with limited clinical relevance. Here, we investigated the cost of a *bla*_CTX-M-1_-IncI1/ST3 epidemic plasmid in a commensal *E. coli* animal strain, before and after oral inoculation of pigs. In our model, *bla*_CTX-M-1_-IncI1/ST3 plasmid imposed no measurable burden on the recipient strain after conjugation and we observed no modification of the bacterial growth rate of the *bla*_CTX-M-1_-IncI1 plasmid-carrying strain after the longitudinal carriage in the pig’s gut. However, a significant difference of bacterial growth rate was found between the *bla*_CTX-M-1_-IncI1/ST3 plasmid-carrying strain and the transconjugants that had spontaneously lost the IncI1 plasmid during *in vivo* carriage, suggesting a small fitness burden exists. The loss of IncI plasmid is a rare event (10), as expected because of the presence of addiction systems on these large conjugative multiresistance plasmids. As a whole, our observations could suggest that *bla*_CTX-M-1_-IncI1 epidemic plasmids do not induce a sufficient fitness cost on its host bacterium to drive their extinction in gut microbiota, in the absence of antibiotics.

After ESBL-producer *E. coli* inoculation, *bla*_CTX-M-1_-IncI1/ST3 plasmid disseminated rapidly to natural isolates in the pig’s gut, highlighting the usually very efficient conjugation system of IncI plasmids (8, 30). As reference the bacterial growth rate of the *bla*_CTX-M-1_-IncI donor strain, the comparative growth rate of the eleven newly formed transconjugants in the pig’s gut appeared extremely variable, related to variable fitness of these IncI plasmid-carrying new spontaneous transconjugants. Although it is not possible to determine the competitiveness of the host cells before IncI plasmid acquisition due to the experimental design of our study, this suggests that the fitness cost could not act as a strong barrier to transmission of the *bla*_CTX-M-1_-IncI1 epidemic plasmid, among natural populations of *E. coli* in a relevant environmental niche such as the gut. Interestingly, the relative abundance of some of the newly formed transconjugants was greater than 50%, suggesting that these clones represented dominant ESC-resistant *E. coli* strains in the pig’s gut at this time.

Previous studies have reported *in vivo* transfers of multidrug-resistant plasmids despite competition from fecal flora, and a high risk of rapid increase of transconjugants becoming the dominant population under antibiotic treatment (30–32). Some studies have shown that antibiotics exposure increased conjugative transfer of *bla*_CTX-M-1_-IncI plasmids (33) and it has been suggested that the transfer rate of the IncI plasmid may be underestimated in *in vitro* study models (34). The high predominance and worldwide spread of some IncI1 plasmids, especially among animal ESBL-producer *E. coli*, could be explained by a low fitness cost imposed on the new recipient strain, combined with a high conjugation efficiency and potentiated by a high exposure of animals to antibiotics in agricultural settings.

A complex plasmid-chromosome cross-talk is involved in plasmid domestication into their host cells and often appears to be dependent of the plasmid-host pairing studied. Among these plasmid-host adaptation effects associated with a potential fitness benefit, only changes in chromosomal gene expression have been observed in some plasmid-host evolutions (35). These modifications in the expression of chromosomal genes for the bacterium adaptation to the acquisition of the plasmid appeared highly variable both in the degree of alteration and in the range of cellular functions that are affected (35–40). However, chromosome-encoded functions most commonly targeted by plasmid carriage include respiration, signaling, energy production, various metabolism pathways, and virulence factors, especially adhesion-related functions involved in biofilm formation and mobility (41, 42). To analyse the impact of the *bla*_CTX-M-1_-IncI1/ST3 plasmid on the commensal *E. coli* host, we performed WGS and RNA-sequencing of the ancestral strains on the one hand, and UB12 isolates collected from pigs’ feces on the other hand. In our study, a very small number of SNPs in chromosomal and plasmid coding sequences were detected in the UB12 isolates, without clear convergence. In addition, RNA-sequencing revealed that only the expression of a small set of chromosomal genes was altered in the host cell after acquisition of the IncI ST3 plasmid, involving chromosomal genes encoding prophages proteins and genes found in different processes such as metabolism, stress response, and cell mobility. We showed that the transcriptional expression of chemotaxis and flagellar genes change between IncI plasmid-carrying and plasmid-free UB12 isolates collected after oral inoculation of pigs, in accordance with the modification in the swimming capacities of these clones. None of mutations observed in our data were associated with change in gene expression, suggesting that we observed a physiological response. It is interesting to note that chromosomal genes implicated in cell mobility are also involved in the bacterial response to environmental changes especially in the highly dynamic and compartmentalized gut environment (43). In our model, plasmid-host adaptation could be the result of both the specific plasmid-host pairing studied and adaptation to environmental changes in the pig’s gut. Overall, we observed that the *bla*_CTX-M-1_-IncI1/ST3 plasmid has only a limited effect on the host cell, targeting cellular functions classically known to be affected in the host bacterium after plasmid acquisition.

We observed that most of the *bla*_CTX-M-1_-IncI1/ST3 plasmid genes were expressed at low level or not expressed in host cell. Our observation indicates that the IncI plasmid transcription is highly regulated in this model, which is consistent with findings from other studies reporting transcriptional profiles of other large conjugative plasmids (39). Recent studies have reported the involvement of plasmid genes encoding for nucleoid-associated proteins (NAPs) homologs, such as H-NS-like proteins, in transcriptional regulation networks between plasmids and host chromosome (42, 44–46). Plasmids encoding H-NS-like “stealth” protein reduce their fitness cost by silencing transcriptional activities of their own genes, as well as some chromosomal genes, and therefore minimize the impact of plasmid introduction in the bacterium. Large conjugative plasmids frequently carry NAP gene homologs but, using *in silico* analysis, similar genes were not yet identified in plasmids of the IncI incompatibility group (47). However, a large number of IncI backbone genes are yet annotated as hypothetical proteins. Future studies could be interesting to characterize new regulators still unknown among these genes, and to better understand the cross-talk between IncI plasmids and the host chromosome.

Our study has further limitations. First, we performed the experiments in a unique commensal *E. coli* host strain and the genetic background of strains plays an important role in plasmid-host adaptation. Second, due to the design of our study, we used *in vitro* plating of the bacteria and therefore we could not directly observe the *E. coli* populations of interest in the pig’s gut. This approach might have introduced a bias as a consequence of short adaptation of the IncI plasmid in the new recipient strain. Nevertheless, our study is the first to provide direct evidence of *in vivo* plasmid transfer in the pig’s gut and to confirm that *bla*_CTX-M-1_-IncI1/ST3 plasmid fitness cost on its bacterial hosts is negligible *in vivo*, in a relevant environmental niche such as the gut.

## MATERIALS AND METHODS

### Bacterial strains and plasmids

For this study, a total of 24 strains were selected from a pig strain collection of ANSES laboratory (Ploufragan, France). The *E. coli* transconjugant UB12-TC was obtained by conjugation between a randomly chosen pan-susceptible pig commensal *E. coli* isolate, made resistant to 250 mg/liter rifampicin (recipient strain UB12-RC, B1, O100:H40) and an ESBL-resistant *E. coli* pig isolate (donor strain M63-DN, B1, O147:H7). Each strain belongs to ST2520 and ST2521, respectively, according to the Warwick MLST scheme.

Thus, *E. coli* UB12-TC is resistant to rifampicin and harbors a IncI1 pST3 plasmid encoding *bla*_CTX-M-1_, *sul2, dfrA17*, and *aadA5* genes. Seven cefotaxime- and rifampicin-resistant isolates (named UB12-TC^ive^), three cefotaxime-susceptible and rifampicin-resistant isolates (named UB12-TCΔpIncI), and eleven cefotaxime-resistant and rifampicin-susceptible isolates (named NI-1 to NI-11) were chosen among all strains collected during a series of experiments conducte to evaluate the carriage of ESBL-producing *E coli* in the pig’s gut (11, 12). These isolates have been collected from day 4 to day 45 after pig inoculation.

### Animals and experimental design

As previously described in Mourand *et al*. (10, 12), the experiments were conducted in the ANSES, Ploufragan, France, animal facilities. The trials were conducted with 6-to 8-week-old specific pathogen-free and initially ESC-resistant *E. coli-* free Large White piglets. The animals were randomized before the experiment. They did not receive any antibiotic treatment prior to the trial and were given the same non-supplemented feed. Each room contained eight pigs placed in two pens of four animals each. For both trials, UB12-TC was inoculated on the first day (day 0): each pig was orally given a 10-ml suspension prepared from strain cultivated on MH agar containing cefotaxime (2 mg/liter). Individual fecal samples were collected from animals on day 0 before inoculation, and between day 1 and 45. The fecal samples were stored at -70°C until analysis.

The numbers of cefotaxime-resistant *E. coli* (including UB12-TC and NI) in the individual fecal samples were estimated by spreading 100 μl of 10-fold dilutions on MacConkey agar plates (containing respectively 2 mg/liter cefotaxime), in triplicate. The titers of total *E. coli* population were determined by plating serial dilutions on MacConkey agar plates without antibiotic. After incubation at 37°C, the colonies obtained on supplemented or non-supplemented MacConkey agar plates were enumerated, and the titers were calculated for each pig per day. The relative abundance of cefotaxime-and rifampicin-susceptible (NI clones) was calculated based on the ability of 2 to 5 isolates obtained from each pig on each day on cefotaxime-supplemented media to grow on rifampicin-supplemented media, and expressed as a percentage.

### Antimicrobial susceptibility testing

Antimicrobial susceptibility testing by the disk diffusion method (ampicillin, ticarcillin, piperacillin, amoxicillin-clavulanic acid, ticarcillin-clavulanic acid, piperacillin-tazobactam, cefotaxime, ceftazidime, aztreonam, cefoxitin, cefepime, ertapenem and imipenem, association of sulphonamides + trimethoprim, tetracycline, streptomycin, nalidixic acid, ciprofloxacin, and rifampicin) was performed according to European Committee on Antimicrobial Susceptibility Testing (EUCAST) guidelines (http://www.eucast.org).

### Motility assay

Donor strain M63-DN, recipient strain UB12-RC, ancestral UB12-TC, all UB12-TC^ive^ and UB12-TCΔpIncI clones were grown overnight in LB medium, washed, suspended in isotonic water to a concentration of 1 ×10^2^ cells ml^-1^ and used to inoculate semi-solid LB agar tubes containing 0.35% agar, incubated for 48 h at 37°C. Experiments were repeated 3 times. For data analysis, three categories were considered (motile, partially motile, and nonmotile).

### In vitro *evolution experiments*

Five randomly selected colonies of UB12-RC and UB12-TC were inoculated into 5 ml of LB and grown under constant shaking at 200 rpm. After overnight incubation at 37°C, 5 μl of each culture was inoculated into five tubes containing 5 ml of fresh LB. At this step, the 10 cultures were considered the ancestors of each lineage (day 0 of the evolution assay). Next, the 10 cultures were propagated by daily serial transfer, with a 1/1,000 dilution (5 μl of culture grown overnight diluted into 5 ml of fresh LB without antibiotic), for 30 days (corresponding to 300 generations in total) and incubated overnight at 37°C under constant stirring at 200 rpm. Finally, after 30 days, 10 independent lineages were obtained (denoted “ite”). At day 0 and every 5 or 6 days of the evolution assay, the culture was isolated on LB agar for purity, and antibiotic susceptibility was tested by disk diffusion. A culture sample from each lineage was collected and stored at -80°C with glycerol for future analysis.

### Individual fitness assays

Fitness, defined here as the Maximum Growth Rate (MGR), was determined from growth curves using a high-precision technique developed in-house as described previously (18). For each strain, the fitness assay was repeated six times. The measurements were collected using a Python script developed in our laboratory, and the MGR was obtained by the lm function in R (http://www.R-project.org/).

### Whole-genome sequencing

WGS was performed using Illumina technology. Briefly, DNA was prepared for paired-end sequencing on the Illumina MiniSeq (Integragen, MA, USA) for all strains. DNA samples were extracted using the genomic DNA NucleoMag tissue kit from Macherey-Nagel (Hoerdt, France). DNA libraries were prepared and indexed, and tagmentation was processed using the Nextera DNA Flex library Prep kit (Illumina, USA). The sample concentrations were normalized in equimolar concentration at 1 nM. The pooled libraries were sequenced to a read length of 2 by 150 bp with Illumina MiniSeq High output reagent Kit. The genomes were sequenced at an average depth of 50X.

### DNA sequence analysis and UB12-TC genome comparison

Illumina genome sequence reads were assessed for quality using FastQC v0.11.8 and subsequently trimmed with a cutoff phred score of 30 using TrimGalore v0.4.5. All genomes were *de novo* assembled using SPAdes v.3.13.1 software and analysed with an in-house bioinformatics pipeline (https://github.com/iame-researchCenter/Petanc) adapted from Bourrel et *al*. (48). Briefly, genome typing was performed including MLST determination, phylogrouping, and serotyping. The pipeline was also used to get information about plasmid replicons, resistance and virulence genes, the prediction of a plasmid sequence or a chromosome sequence for each contig using PlaScope (49).

For each sample, sequenced reads were mapped to a reference sequence to analyse the mutations and rearrangements. For reference plasmid sequences, we used the plasmid sequences extracted from GenBank (http://www.ncbi.nlm.nih.gov/GenBank/) (ColRNAI, accession n° J01566; IncY, accession n° MH42254; Col156, accession n° EU090225 and IncI, accession n° KJ484635) and *de novo* assembled contigs as predicted to plasmid genome of UB12-TC strain with PlaScope, included in our pipeline. For reference chromosomal genome, first, we calculated the mash distance between UB12-TC genome sequence and *E. coli* genomes from RefSeq database using Mash v.2.0 and we selected the closest *E. coli* strain assembly ASM190094v1. Then, all predicted chromosomal contigs of the UB12-TC genome were rearranged as scaffold with RAGOUT v2.3 (https://github.com/fenderglass/Ragout)(50) and the reference genome (*E. coli* assembly ASM190094v1). All scaffolds of the UB12-TC strain obtained were annotated on the MicroScope platform (51). Identification of the point mutations and rearrangements between UB12-TC and UB12-TC^ive^ were performed using the open computational pipeline Breseq v0.27.1 (26). All SNPs identified in the ancestor clones that were sequenced were subtracted from the sequences of the evolved clones.

### Transcriptome sequencing

RNA sequencing was performed on the recipient strain UB12-RC, the ancestral transconjugant UB12-TC, two evolved UB12-TC transconjugants (UB12-TC^ive^, detected in pig’ stools at day 28 and 31) and one UB12-TC transconjugant that had lost the plasmid IncI (UB12-TCΔpIncI, detected in pig’ stools at day 20).

For RNA extraction, a total of 10 cultures representing two biological replicates per strain were independently grown. For each strain, 6 μl of an overnight culture was added in to 6 ml of fresh LB and was incubated at 37 ° C under constant stirring at 200 rpm until reaching an OD600 of approximately 0.6. 3 ml of each culture were taken, representing two technical replicates, and mixed with RNA protect (Qiagen, Courtaboeuf, France) according to the manufacturer’s recommendations. Cells were immediately pelleted by a 10 min centrifugation at 5,000 g. RNA was prepared using the Quick-RNA Fungal/Bacterial MiniPrep kit (Ozyme, ZymoResearch, USA). DNA was digested using Turbo DNAse (Thermofisher, UK) based on the manufacturer’s protocol. Verification of complete removal of any contaminating DNA was performed by PCR amplification of *recA* housekeeping gene. The final RNA solution was quantified with the Qubit™ RNA High Sensitivity assay kit (Thermofisher Scientific, UK) and RNA quality was evaluated using the Agilent 2100 Bioanalyzer RNA 6000Nano Kit (Agilent Technologies, Santa Clara, CA). The RNA was immediately stored at -80°C.

For each replicate, 8 μl of RNA was treated to remove 16S and 23S rRNAs and cDNA libraries were prepared using the Zymo-Seq RiboFree® Total RNA Library Kit (Ozyme, ZymoResearch, USA). The cDNA quantity was determined with the Qubit™ dsDNA Broad Range assay kit. Paired-end sequencing of the pooled libraries was performed with an Illumina MiniSeq Mid Output reagent kit and an Illumina NextSeq 500/550 High Output Kit v2.5 (Illumina, USA), configured to 2 × 150-bp cycles, yielding approximately 8 and 400 million reads, respectively.

### Transcriptome analysis

To assure high sequence quality, the Illumina cDNA reads in FASTQ format were trimmed to remove all sequencing adapters using TrimGalore v0.4.5., according to the manufacturer’s recommendations. Then, cDNA reads were trimmed again, using TrimGalore v.0.4.5., with a cutoff phred of 30, and the reads shorter than 70 bp were discarded. A final quality check was done using FastQC v0.11.8. Reads were then mapped to the UB12-TC reference genome sequence (see above) using BWA alignment software and BWA-MEM algorithm v.0.7.17. (52) Mapping statistics were verified using BAMStat v.1.25. and SAMtools idxstat v.1.9. (53). Reads counts were performed using featureCounts v.2.0.3. (http://subread.sourceforge.net) (54). An R package, DESseq2 v.1.34. was used to normalize and to identify differentially expressed genes between each sample (http://www.bioconductor.org/packages/release/bioc/html/DESeq2.html) (55). A cutoff of q value of < 0.05 and a fold change of > 2 were used to measure statistical significance.

### Statistics

For statistical data analysis from the fitness assays, we used an analysis of variance (ANOVA) and unpaired Student’s test using R software, with *P* values of < 0.05 being considered significant.

### Ethics

The experiments were performed in accordance with French animal welfare regulations, and the protocol was approved by the ANSES/ENVA/UPEC ethical committee (ComEth authorizations 12-032 and 15-050 [no. APAFIS: 2015061813246553]). The experiments were conducted in the ANSES, Ploufragan, France, animal facilities.

### Data availability

All reads have been deposited at the European Nucleotide Archive (project accession number PRJEB8070, ID samples from ERS14303872 to ERS14303895)

## ACKNOWLEDGMENTS

We thank Jean Marc Guigo (Institut Pasteur) for useful discussions and Olivier Clermont, Kevin La and Gwenaelle Mourand (ANSES) for technical help.

This work was partially supported by a grant from the Fondation pour la Recherche Médicale to E.D. (grant number DEQ20161136698), ANSES (French Agency for Food, Environmental, and Occupational Health and Safety), France-Agrimer (grant 2012-0388), and by the Conseil Départemental des Côtes d’Armor. The funders had no role in study design, data collection and interpretation, or the decision to submit the work for publication.

